# Development of a humanized mouse model to analyze antibodies specific for human leukocyte antigen (HLA)

**DOI:** 10.1101/2020.07.13.200394

**Authors:** Senichiro Yanagawa, Hiroyuki Tahara, Takayuki Shirouzu, Shintaro Kawai, Yuka Tanaka, Kentaro Ide, Shuji Akimoto, Hideki Ohdan

**Affiliations:** Department of Gastroenterological and Transplant Surgery, Graduate School of Biomedical and Health Sciences, Hiroshima University, Hiroshima, Japan; Molecular Diagnostics Division, Wakunaga Pharmaceutical Co., Ltd., Osaka, Japan

## Abstract

In organ transplantation, human leukocyte antigen (HLA)-mismatch grafts not only induce the activation of cellular mediated immune response but also the development of chronic antibody-mediated rejection due to the donor-specific anti-HLA antibody (DSA) produced by B cells and plasma cells interacting with the graft endothelium.

Significant improvement in long-term survival after transplantation can be expected if antibody-mediated rejection due to the DSA can be overcome. However, the mechanism of producing or controlling the DSA remains to be elucidated.

In recent decades, “humanized mouse model” have been widely used for the basic research of human immune systems, but a humanized mouse model to analyze the mechanism of DSA production has not been established yet. Thus, we aimed to create a humanized mouse using a severe immunodeficiency mouse (NSG mouse) administered with human peripheral blood mononuclear cells (PBMCs). Initially, we detected very low level of human total-IgG and no anti-HLA antibodies (Abs) in these mice. The responder PBMCs with antibody-producing B cell activating factors added or regulatory T cells depleted were subsequently co-cultured with the irradiated stimulator PBMCs *in vitro*, and these whole cells were administered into naïve NSG mice. The humanized model with sufficient human total-IgG and anti-HLA antibody production was consequently established. Interestingly, in all these mouse models, allo-specific anti-HLA Abs production was prominently suppressed, whereas non-allo-specific anti-HLA Abs were sufficiently detectable.

Therefore, this novel humanized mouse model might be useful for analyzing the mechanism of anti-allogeneic human B cell tolerance induction.

## Introduction

Human leukocyte antigen (HLA) is distributed in nearly all cells and body fluids and functions as a histocompatibility antigen (an important molecule related to human immunity). In organ transplantation, HLA conformity is important because different forms of HLA are recognized as foreign objects which are subject to attack from the immune system; additionally, HLA-mismatched grafts induce the activation of cellular mediated immune response, leading to graft rejection in the absence of immunosuppressive therapy [1-4]. Therefore, appropriate selection of donors is required before transplantation. However, HLA is rich in polymorphisms, and HLA between recipients and donors differ in many cases. Despite recent advances in immunosuppression and antibody (Ab) screening prior to transplantation, chronic antibody-mediated allograft rejection due to donor-specific anti-HLA antibodies (DSA) influence transplantation outcomes; however, the immunological mechanism of antibody-mediated rejection due to the DSA is unclear [1,2]. Elucidation of the antibody-mediated rejection is imperative for developing new immune therapies to improve long term prognosis of organ transplants [5]. To develop a therapy that effectively regulates DSA-secreting cells, it is necessary to determine the mechanism of the DSA production in human immunocompetent cells.

In mouse models of heart transplantation from Balb/c to B6, the spleen and bone marrow are the major sources of DSA-secreting cells [6]. However, the detection of DSA-secreting cells and mechanism of DSA production in *in vivo* human immune cells have not been widely examined because *in vivo* analysis of the DSA production in human clinical setting is difficult. Successful immunotherapies produced in animal models and transplanted into clinical cases have shown limited success because of the many species-specific differences between mouse and human immune responses.

In recent decades, immunodeficient mice for engraftment with the functional human immune system have been developed, known as “humanized mouse models” [7-11]. These models contain various types of human cells and tissues engrafted in immunodeficient mice and are extremely useful for basic research, particularly for studies of the human immune system. However, there is no established humanized mouse model that analyzes the mechanism of human DSA production and the antibodies (Abs)-secreting human B cells, especially one that uses human peripheral blood mononuclear cells (PBMCs).

To detect human DSA-secreting cells and the DSA in an *in vivo* model, we attempted the establishment of an anti-HLA Ab producing humanized mouse model by reconstructing human immunocompetent cells. First, we have tried to establish the humanized mouse using PBMCs as below: two types of PBMCs with different HLA types (Table1; combination 1) were administered to severe immunodeficient naïve NSG mouse to produce anti-HLA Abs. Responder PBMCs were intravenously injected to naïve NSG mice (Mouse # S-1 and S-2), then 1 day later, the mice was challenged with irradiated stimulator PBMCs. In FCM, only the responder cells were detected in the peripheral blood of humanized mouse (S1 Fig.). However, we detected very low level of human total-IgG and no anti-HLA Abs in these mice (S2 and S3 Figs.). Responder PBMCs with Ab-producing B cell activating factors added or regulatory T cells depleted were *in vitro* were co-cultured with the irradiated stimulator PBMCs before administration into naïve NSG mouse in order to facilitate anti-HLA Ab production.

## Materials and methods

### Ethics statement

This study was performed in strict accordance with the Guide for the Care and Use of Laboratory Animals and the local committee for experiments. The experiment protocol was approved by the Ethics Review Committee for Animal Experimentation of the Graduate School of Biomedical Sciences, Hiroshima University (Permit Number: A17-64-2). All animal experiments were performed according to the guidelines established by the US National Institutes of Health (1996). This work was carried out, in part, at the Research Facilities for Laboratory Animal Science, Natural Science Center for Basic Research and Development, Hiroshima University.

### Mice

NOD.Cg-Prkdc^scid^IL2rg^tm1Wjl^/SzJ (NSG) mice, aged 6 weeks, were purchased from Japan Charles River (Yokohama, Japan) and maintained and bred in a specific pathogen-free facility in micro isolator cages at the Natural Science Center for Basic Research and Development Hiroshima University. Animal welfare was carefully ensured by experienced operators every day. All efforts were made to minimize the suffering of animals for the duration of their lives.

### Combination of responder and stimulator’s HLA alleles

Three different combinations of HLA classI molecules were prepared (Table 1).

**Table 1.** Combination HLA of Responder and Stimulator PBMCs.

| Combination 1 | Combination 2 | Combination 3 |
| --- | --- | --- |
| Responder HLA | Responder HLA | Responder HLA |
| A*24:02, A*33:03 | A*02:01, A*02:06 | A*02:01, - |
| B*15:18, B*52:01 | B*51:01, - | B*13:01, B*40:01 |
| C*01:02, C*12:02 | C*08:01, C*14:02 | C*03:04, C*15:02 |
| Stimulator HLA | Stimulator HLA | Stimulator HLA |
| A*02:07, A*26:01 | A*24:02, A*33:03 | A*24:02, - |
| B*40:02, B*51:01 | B*15:18, B*52:01 | B*51:01, B*54:01 |
| C*03:04, C*14:02 | C*01:02, C*12:02 | C*01:02, C*14:02 |

PBMCs of these combinations were obtained from human recipients or living donors of liver transplant and kidney transplant at Hiroshima University Hospital. Flow cytometry (FCM) analyses were performed to distinguish the cells based on differences in HLA-A2 or A24. All recipients and donors were typed for HLA-A, -B, and -C at the HLA Foundation Laboratory, Kyoto in Japan or using commercially available DNA typing reagent, WAKFlow^R^ HLA DNA Typing (Wakunaga Pharmaceutical Co., Ltd., Osaka, Japan).

### Preparation of feeder cells

Dr. Freeman GJ kindly provided NIH 3T3 fibroblasts expressing the human CD40 ligand (h-CD40L), which is resistant to G418 [12]. This is the basic feeder cells and human B-cell activating factor from the tumor necrosis factor family (h-BAFF) complementary DNA in the pCMV3-untagged vector was transfected into these feeder cells, after which hygromycin-resistant stable clones were selected. The new feeder cells were expressing h-CD40L and h-BAFF. In FCM analysis, the h-BAFF expression rate was over 90 % (S4 Fig.).

Both feeder cells were irradiated with 120Gy gamma radiation before culturing.

### Depletion of regulatory T cells (T-regs) defined CD4+CD25+

Regulatory T cells (T-regs) were depleted that CD4^+^ T cells were negative selection and CD25+ cells were positive selection using the Easy Sep™ Human CD4^+^CD25^+^ T Cell Isolation Kit (StemCell Technologies, Vancouver, Canada). We confirmed the removal of about 90 % of T-regs by FCM (S5. Fig.).

### Generating a humanized mouse model

Three protocols were employed in attempt to generate the humanized model.

#### The first protocol

Two different HLA types of human PBMCs (fresh non-irradiated responder and irradiated stimulator PBMCs) were co-cultured with feeder cells expressing h-CD40L for three days in RPMI-1640 medium supplemented with 10% fetal bovine serum, 5.5 × 10^−5^ M 2-ME, and 10 mM HEPES in a 6-well plate. Mice # 1-1-1 and 1-1-2 were in combination 1, # 1-2-1 and 1-2-2 were in combination 2, # 1-3-1 and 1-3-2 were in combination 3.

#### The second protocol

Two different HLA types of human PBMCs (fresh non-irradiated responder and irradiated stimulator PBMCs) were co-cultured with feeder cells expressing both h-CD40L and h-BAFF for three days in the same condition. Mice # 2-1-1 and 2-1-2 were in combination 1, # 2-2-1 and 2-2-2 were in combination 2, # 2-3-1 and 2-3-2 were in combination 3.

#### The third protocol

Two different HLA types of human PBMCs (fresh non-irradiated T-regs-depleted responder and irradiated stimulator PBMCs) were co-cultured with feeder cells expressing h-CD40L for three days in the same condition. Mice # 3-1-1 and 3-1-2 were in combination 1, # 3-2-1 and 3-2-2 were in combination 2, # 3-3-1 and 3-3-2 were in combination 3.

Table 2 summarizes the protocols as mentioned, *in vitro* co-culture conditions to generate the three different types of humanized mouse model, and the identification number of each experimental mouse.

**Table 2.**
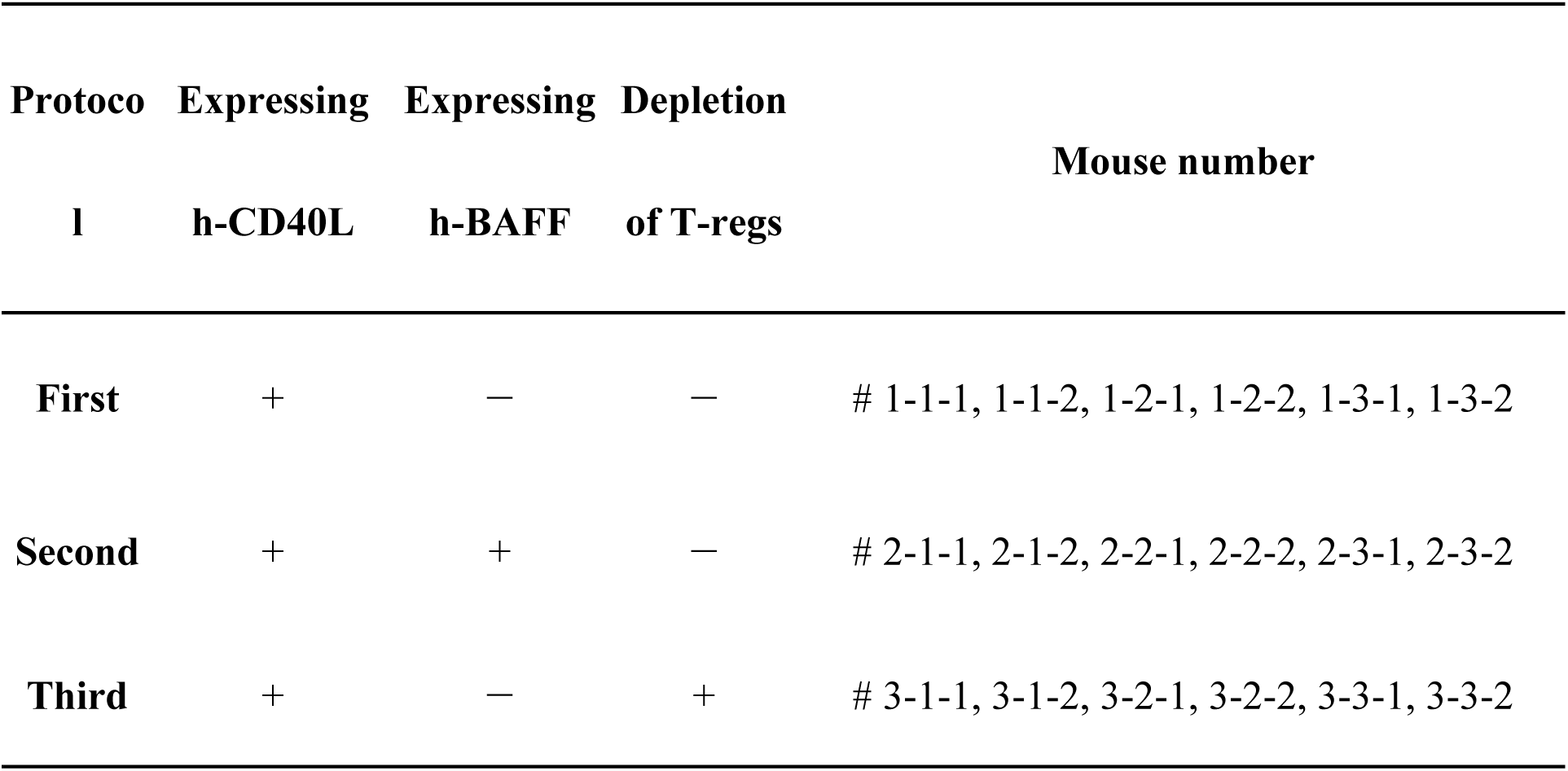
Humanized mouse protocols and pre-culture conditions.

After co-culture in a humidified atmosphere at 37°C with 5% CO_2_ for three days, *in vitro* cultured cells including feeder cells, were collected and directly injected into NSG mouse spleen. On the next day, mice were challenged with intraperitoneal administration of the same stimulator PBMCs with 30 Gy gamma radiation. All experiments were conducted using feeder cells not exceeding 60–70% confluence to avoid pericellular hypoxia induced by over confluency. The cells were collected and counted by the trypan blue dye exclusion method.

### Analysis of flow cytometry (FCM)

Cell suspensions were prepared from peripheral blood of humanized mice at 2 weeks post-injection (day 14) and the chimera state was confirmed. These samples were stained with fluorescein isothiocyanate (FITC)-conjugated anti-HLA-A2 and anti-human CD257(BAFF), phycoerythrin (PE)-conjugated anti-human CD25, CD45, allophycocyanin (APC)-conjugated anti-human CD3, CD4, CD14, CD19 Abs purchased from eBioscience (San Diego, CA, USA), Medical & Biological Laboratories Co., Ltd. (Nagoya, Japan) and BD Pharmingen (San Diego, CA, USA). Flow cytometry was carried out on a FACS Canto flow cytometer (Becton Dickinson, Mountain View, CA) using FlowJo™_v10.6.2 software (Tree Star Inc., Ashland, OR, USA). Dead cells, identified by light scatter and propidium iodide staining, were excluded from the analysis

### Measurement of human total-IgG and anti-HLA antibody titer

To measure human total-IgG in the sera of humanized mice at pre-injection and 1–5 weeks (day 7 to 35) post-injection, we used the Cytometric Bead Array Flex Set and FCAP Array™ v3.0 Software (BD Biosciences, Franklin Lakes, NJ, USA). Anti-HLA Abs are reported as the mean fluorescent intensity (MFI) in the sera of highest human total-IgG; MFI 100 or more was judged to be Ab titer-positive, measured by the Luminex assay using WAKFlow^R^ class I antibody specificity identification reagent (Wakunaga Pharmaceutical Co., Ltd., Japan).

## Results

### Human total-IgG and anti-HLA Abs are sharply increased in the first protocol humanized mouse model, but no allo-specific anti-HLA Abs are detected

We reconstructed a humanized mouse model in which antibody-producing B cells were pre-activated *in vitro* and then transferred to naïve NSG mice. To activate the Ab-producing B cell using the signaling of human CD40-CD40 ligand, two different HLA types of PBMCs (fresh non-irradiated responder and irradiated stimulator PBMCs) were co-cultured with the feeder cells expressing h-CD40L for 3 days in all combination. The collected whole cells (20 × 10^6^) were administered to naïve NSG mice by intrasplenic infusion (day 0), and then the same stimulator PBMCs with 30 Gy gamma-irradiation (10 × 10^6^) were challenged to the NSG mice (day 1). With regards to FCM on day 14, only the responder cells were detected in the peripheral blood of humanized mice, and most h-CD45^+^ cells were h-CD3^+^ in all combinations (Fig 1). None of the human PBMCs derived from the stimulator were detected because of the effect of gamma-irradiation before the co-culture. This chimeric state was similar in all combinations.

**Fig 1.**
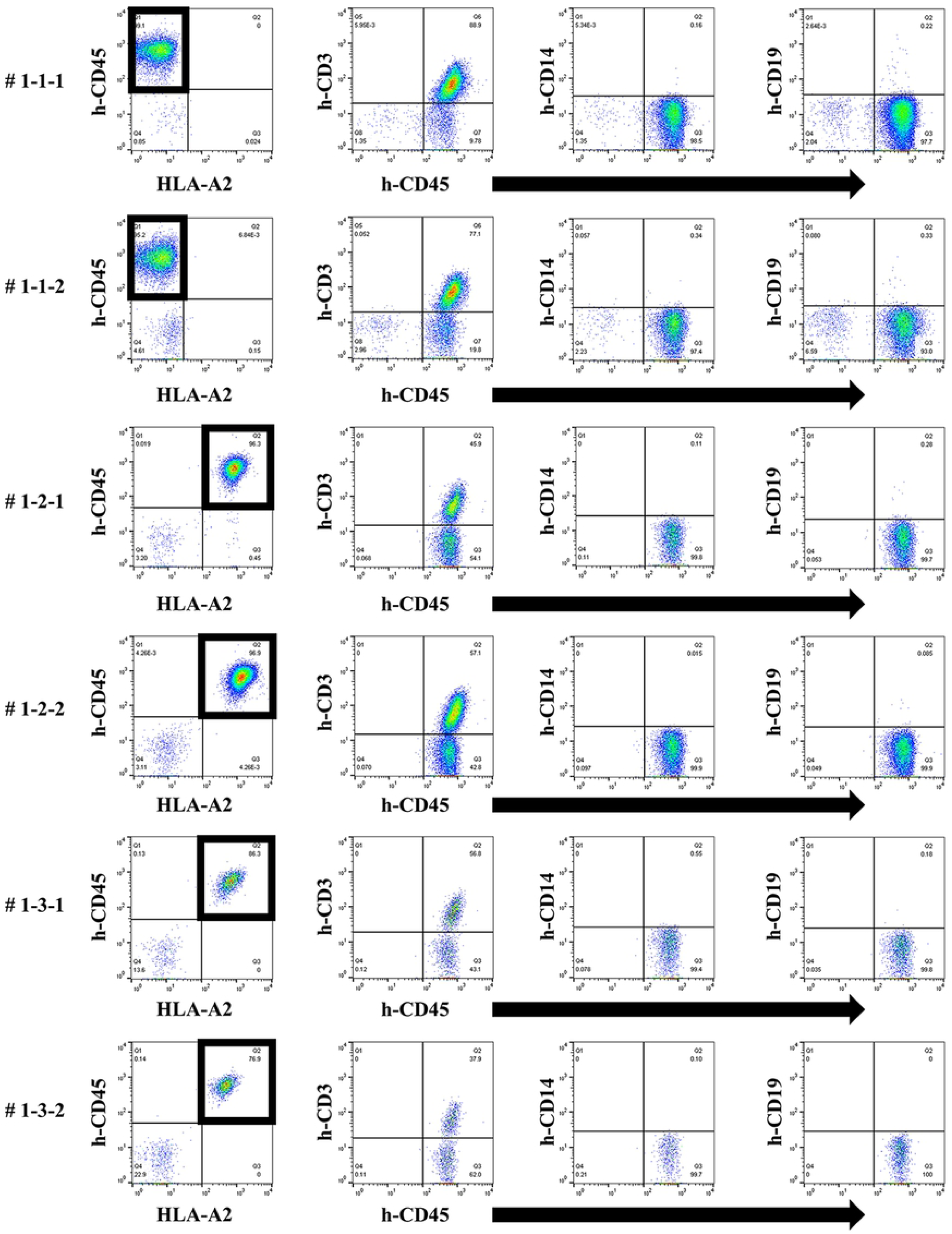
Chimeric status in all humanized mice under the first protocol. In FCM on day 14, only responder cells were detected in the peripheral blood of humanized mice. The cells from the responder are surrounded by square. This chimera status was common in all the humanized mice (# 1-1-1, 1-1-2, 1-2-1, 1-2-2, 1-3-1, 1-3-2). The majority of human CD45^+^ cells were human CD3^+^ T cells and human CD19^+^ B cells were found in all but a few of the mice.

In humanized mouse sera, human total-IgG was increased after administration of the cells to NSG mice (Fig 2). Serum was measured at the point of the highest of human total-IgG and revealed clearly increased anti-HLA Ab titers, but almost all were allo-antigen non-specific anti-HLA Abs in all humanized mice (Fig 3). With each mouse in combination 3, only allo-specific anti-HLA Ab (A*24:02) was slightly produced, but other allo-specific anti-HLA Abs (B*51:01, B*54:01, C*01:02, C*14:02) were not produced completely. No correlation was observed in the titers between human total-IgG and anti-HLA Abs. Moreover, the abundance ratio of peripheral blood B cells in these humanized mice did not show a clear correlation with the values of human total-IgG or anti-HLA Abs. Further, the mice having higher anti-HLA Ab titers did not necessarily show proportionally higher human total-IgG values.

**Fig 2.**
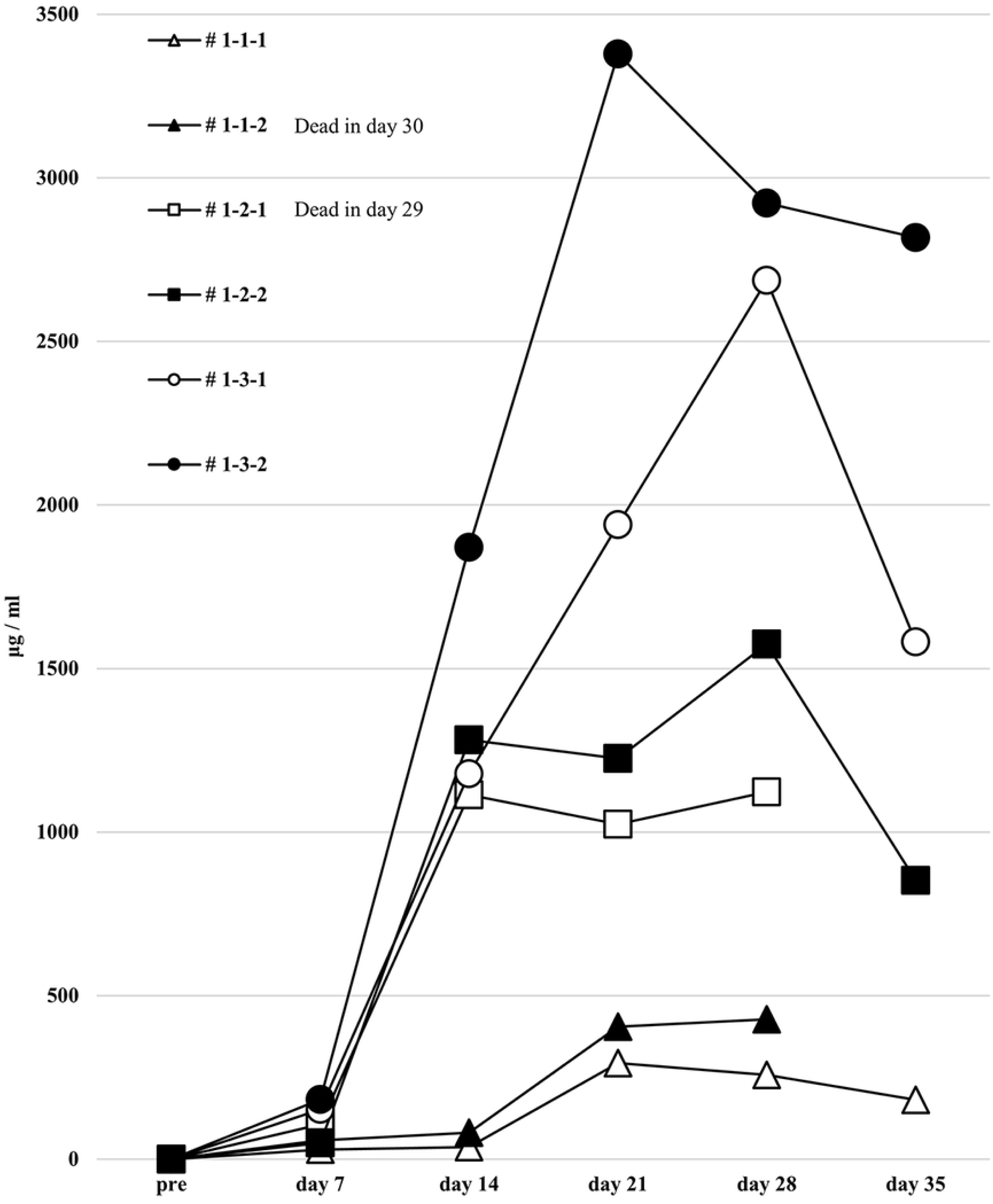
Human total IgG production in all humanized mice under the first protocol. Humanized mouse sera were collected before injection into naïve NSG mice and at 1–5 weeks (day 7 to 35) after injection. Human total-IgG was sufficiently elevated in all the humanized mice. Mouse # 1-1-2 was dead in day 30 and # 1-1-2 was dead in day 29.

**Fig 3.**
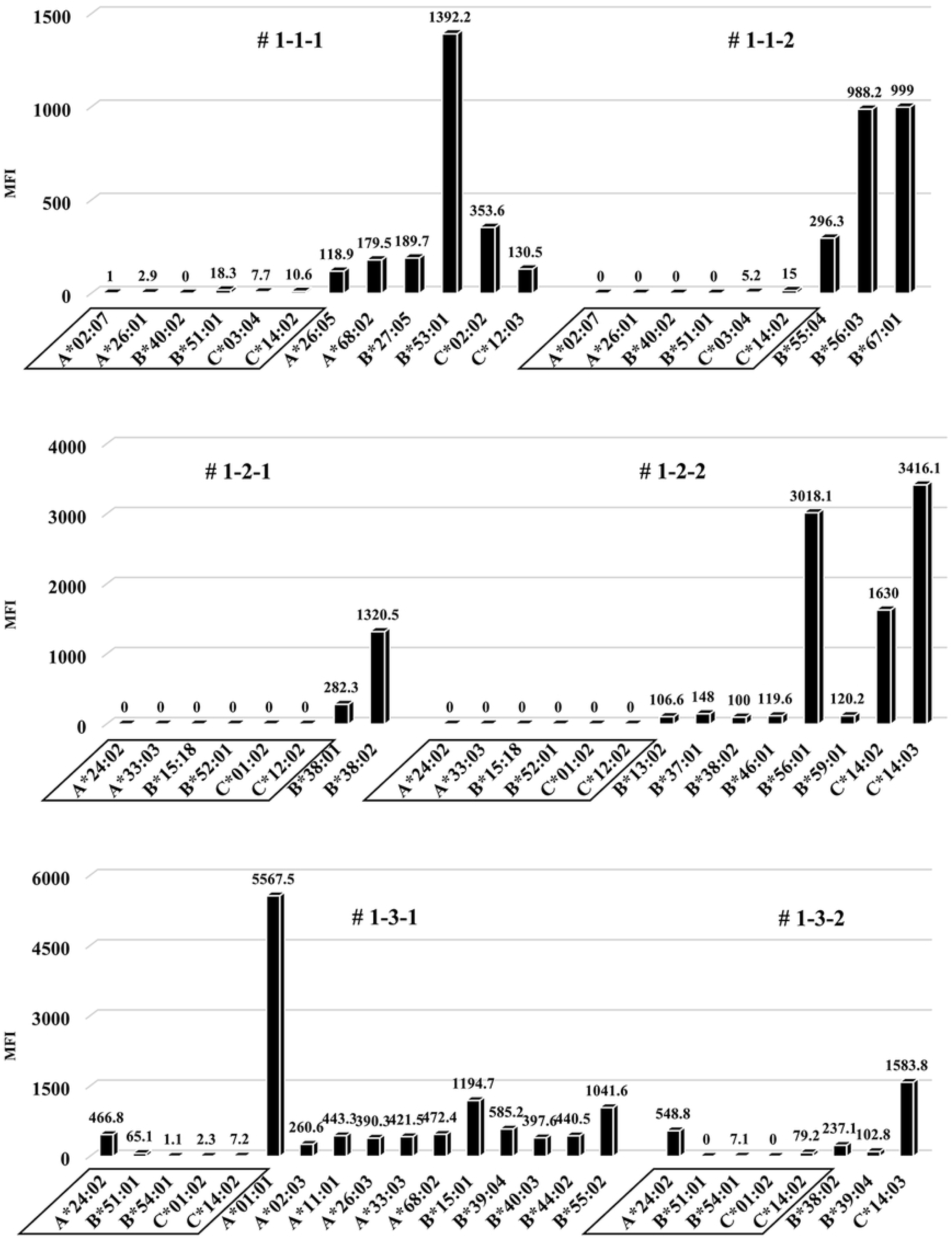
Anti-HLA Abs production in all humanized mice under the first protocol. The results of anti-HLA Abs produced in all of the humanized mice are shown. The stimulator HLA allele name (i.e. allo-specific anti-HLA Abs) are enclosed in a square. All humanized mice produced various allelic non-specific anti-HLA Abs. However, allo-antigen specific anti-HLA Ab (A*24:02) was slightly produced in the mouse # 1-3-1 and # 1-3-2.

### No increase in total human-IgG and whole anti-HLA Abs, and no allo-specific anti-HLA Abs produced in the second protocol of humanized mouse model

To induce better activation of antibody-producing B cells responding to HLA, we utilized the BAFF-BAFF receptor signaling system. Similar experiment as the first protocol was performed by substituting the conditions of feeder cells expressing both h-CD40L and h-BAFF. There were no major changes in the chimera status of the PBMCs from the humanized mice, compared with the first protocol (Fig 4). The results of human total-IgG and anti-HLA Ab were also similar to the first protocol. Further, there was no production of allo-specific anti-HLA Abs, although the total human-IgG production was sufficient (Figs 5 and 6). All the humanized mice had a small amount of the responder CD19^+^ B cells in their peripheral blood (Fig 4).

**Fig 4.**
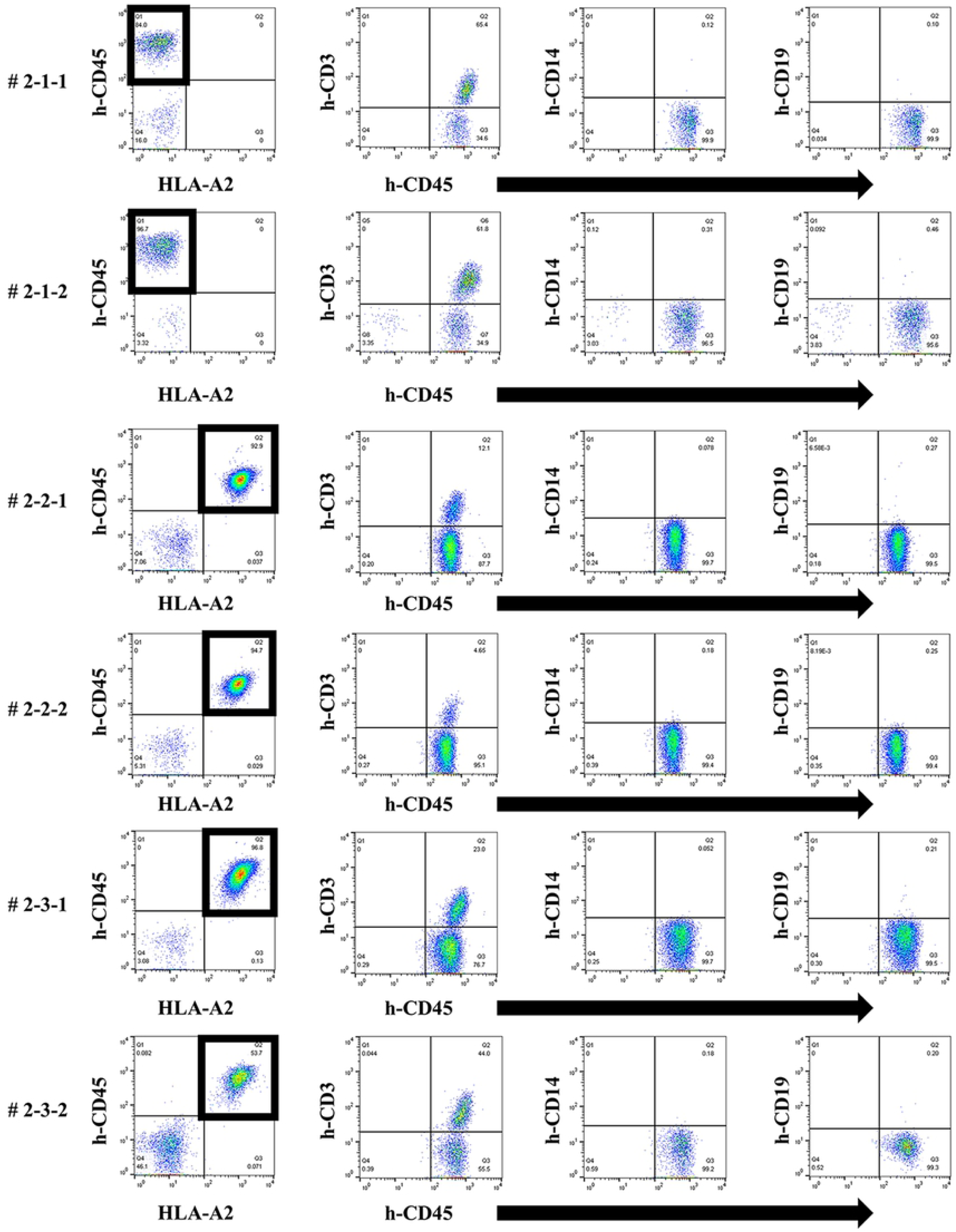
Chimeric status in all humanized mice under the second protocol. By FCM on day 14, only responder cells were detected in the peripheral blood of humanized mice, but not stimulator cells. The cells derived from the responder is surrounded by square. This chimera status showed the same results as the first protocol.

**Fig 5.**
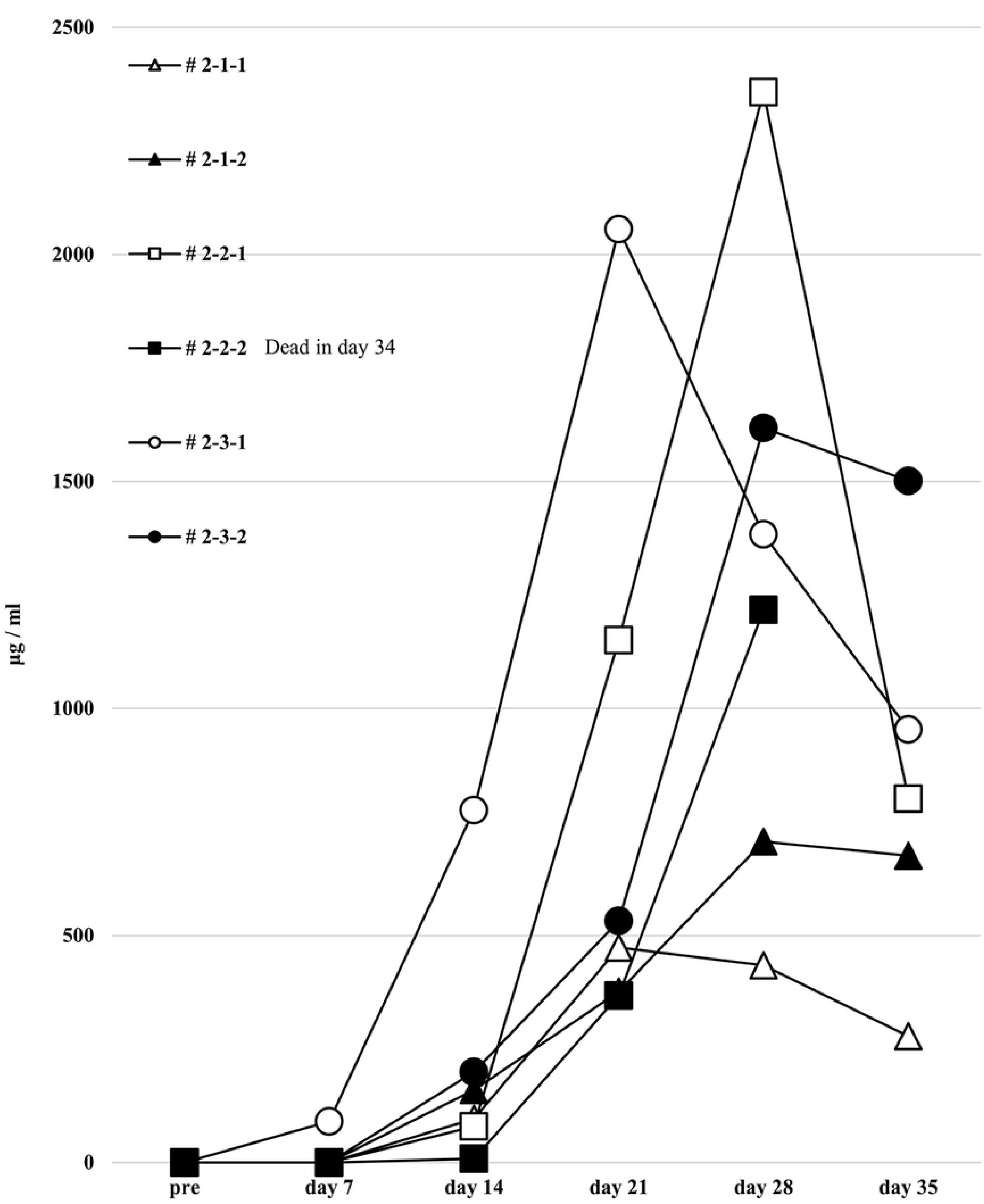
Human total IgG production in all humanized mice under the second protocol. Humanized mouse sera were collected and human total-IgG was measured as described in the first protocol, and the results showed the same tendency as the first protocol. Mouse # 2-2-2 was dead in day 34.

**Fig 6.**
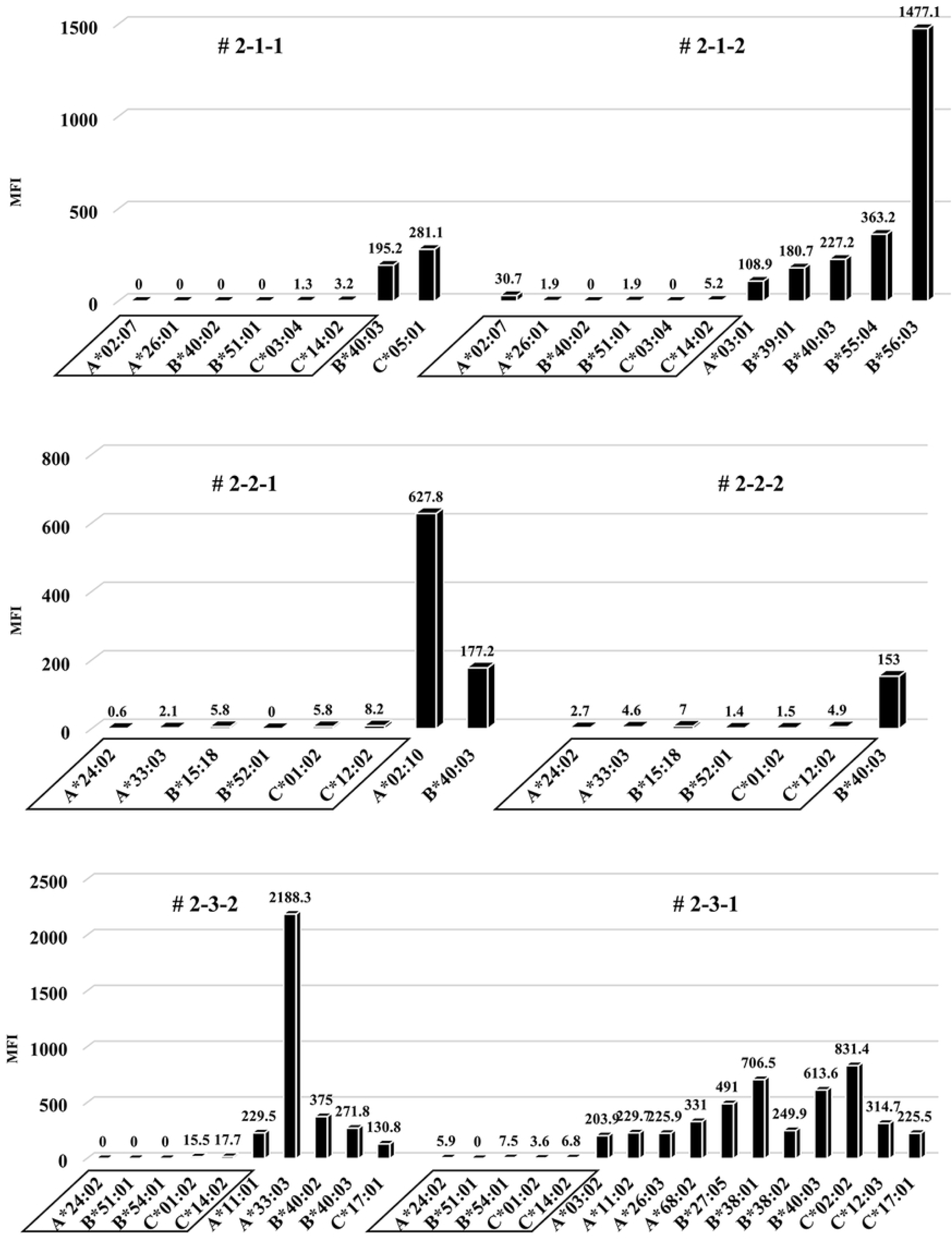
Anti-HLA Abs production in all humanized mice under the second protocol. The results of anti-HLA Abs production in all humanized mice are shown. The stimulator HLA allele name (i.e. allo-specific anti-HLA Abs) are enclosed in a square.

### No remarkable changes of results in total human-IgG, whole anti-HLA Abs production and allo-specific anti-HLA Abs in the third protocol humanized mouse model

Considering that T-regs in the responder’s PBMCs may have interfered with the facilitation of the anti-HLA Ab production during culture, the same protocol as the first protocol was performed under conditions without T-regs—responder PBMCs were co-cultured with the feeder cells expressing h-CD40L. All results, including chimeric status, human total-IgG, and whole anti-HLA Abs, were similar to the first protocol. Although a slight production of allo-specific anti-HLA Abs were observed in the mouse # 3-2-2 and # 3-3-2 (Figs 7-9), however the proportion of responder B cells was not higher in the PBMCs, neither was the human total-IgG titers remarkable.

**Fig 7.**
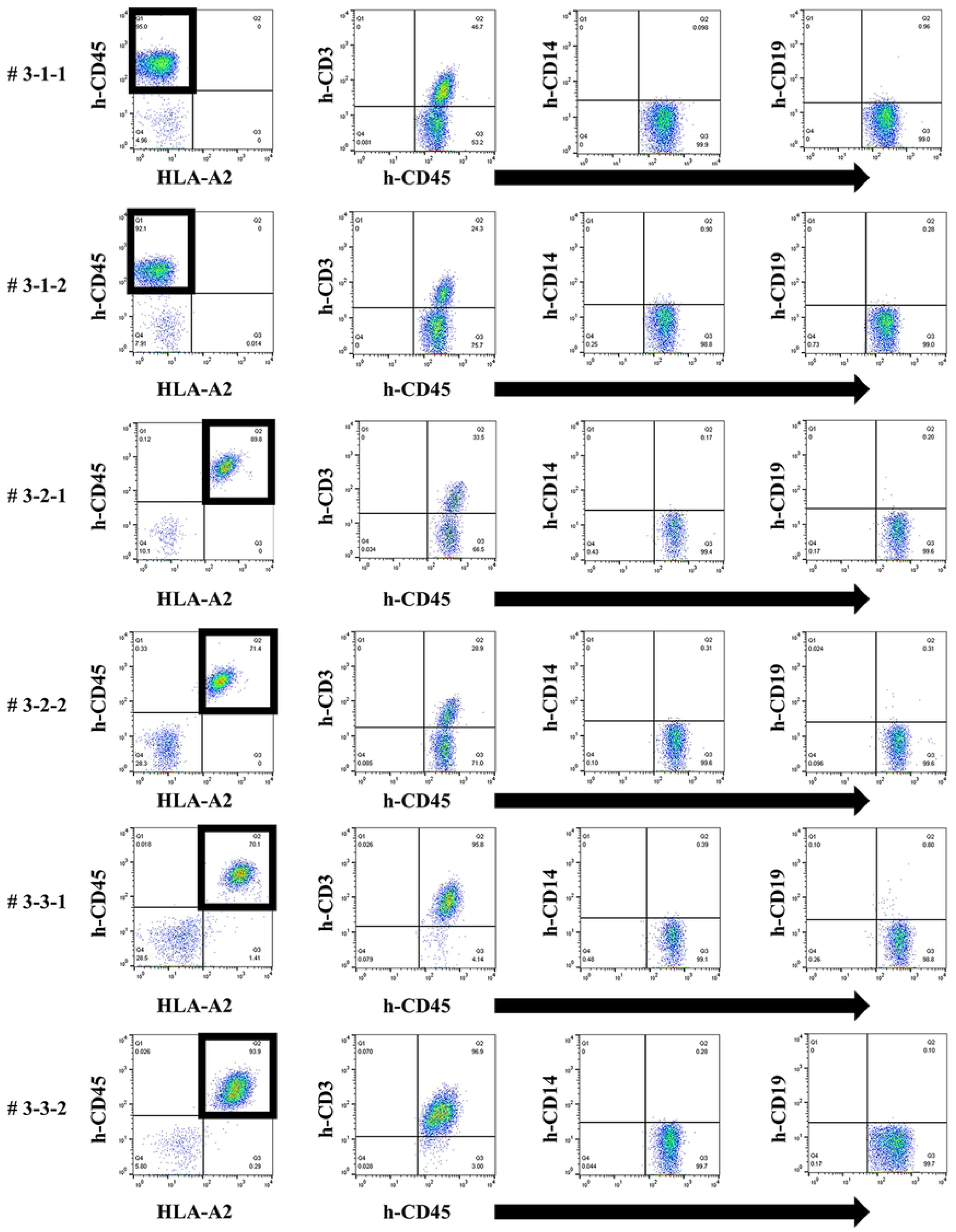
Chimeric status in all humanized mice under the third protocol. By FCM on day 14, only the responder cells were detected in the peripheral blood of humanized mice. The cells derived from the responder is surrounded by square. This chimera status showed the same results as the first protocol.

**Fig 8.**
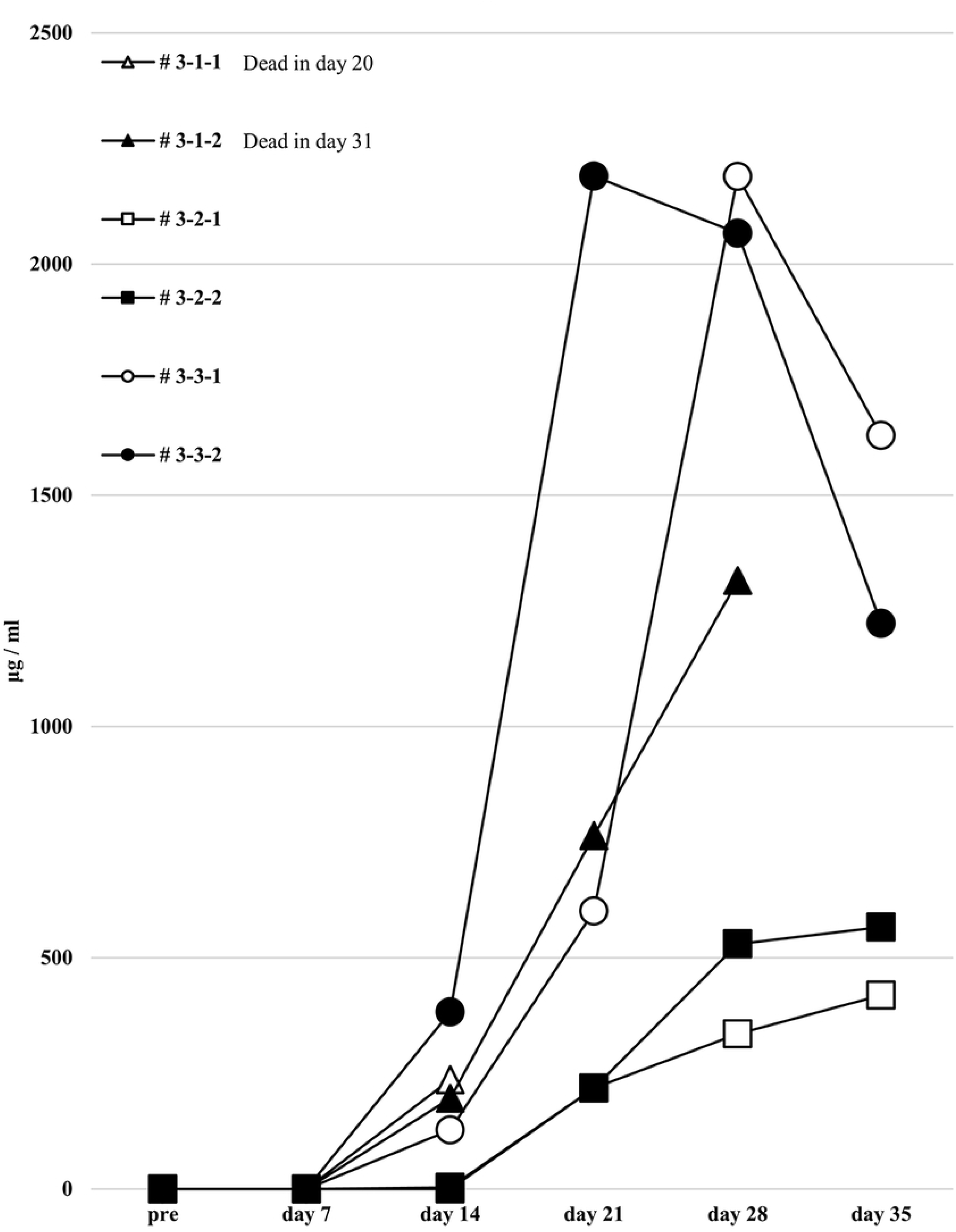
Human total IgG production in all humanized mice under the third protocol. The human total-IgG was measured as described in the first protocol. Human total-IgG titers showed no remarkable changes compared to the first protocol. Mouse # 3-1-1 was dead in day 18 and # 3-1-2 was dead in day 31.

**Fig 9.**
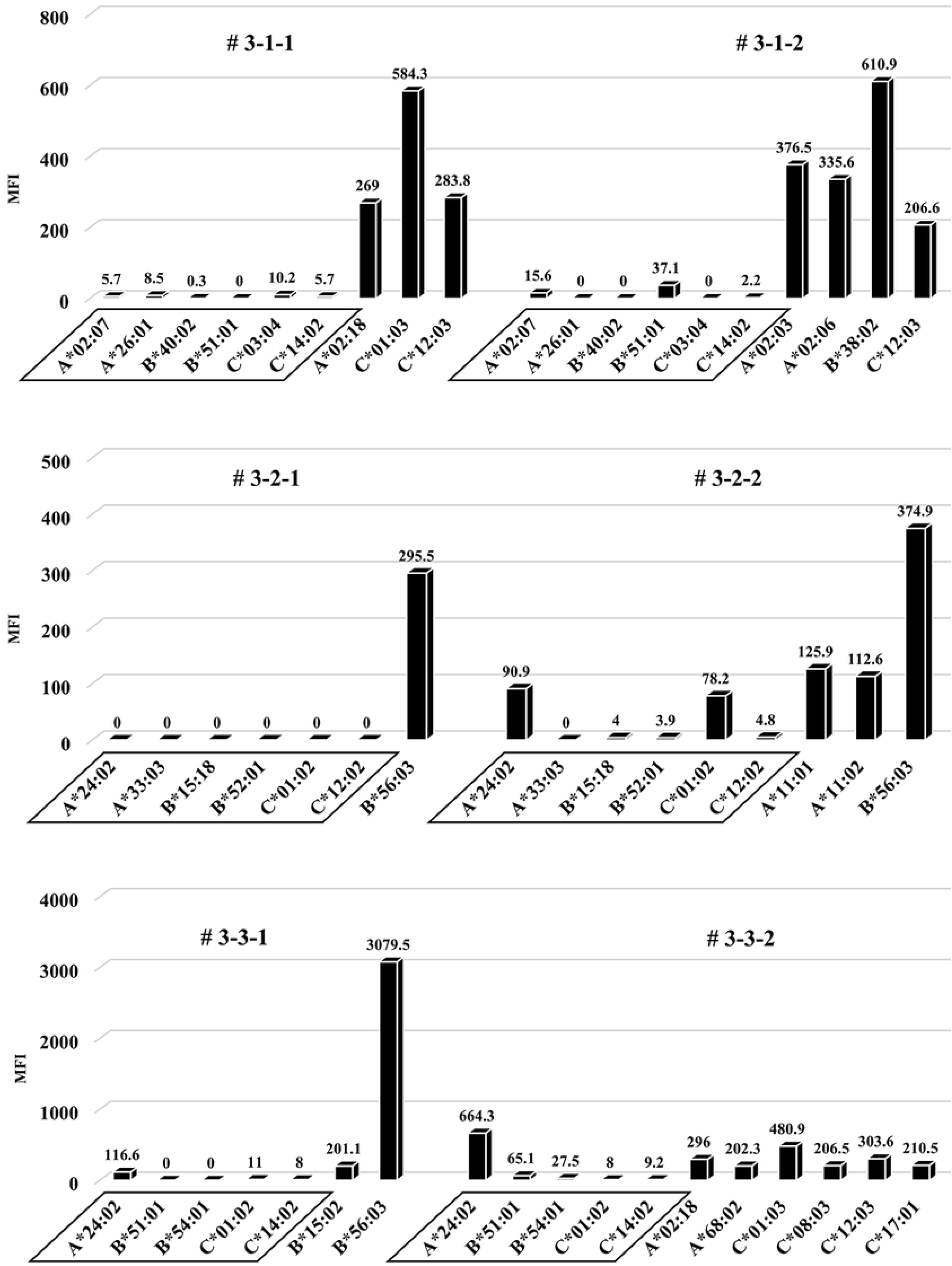
Anti-HLA Abs production in all humanized mice under the third protocol. The results of anti-HLA Abs produced in all of the humanized mice are shown. The stimulator HLA allele name (i.e. allo-specific anti-HLA Abs) are enclosed in a square. All humanized mice have produced various allelic non-specific anti-HLA Abs similar to the result in Fig 3. On the other hand, allo-antigen specific anti-HLA Ab (A*24:02) was slightly produced in the mouse # 3-3-1 and # 3-3-2.

## Discussion

Acute or chronic antibody-mediated rejection involving DSA is one of the most important issues to be overcome in clinical organ transplantation. However, the mechanisms and treatments of antibody-related rejection remain unresolved. The research of human immune responses and disease *in vivo* including anti-HLA Abs production are limited by many constraints. There is need for animal models that on the one hand accurately mirror the pathogenesis of disease, and on the other allow pre-clinical examination of human cell-based therapeutic approaches *in vivo* model.

Moiser *et al*. reported in 1988 that injection of human peripheral blood leukocytes can result in the stable long-term reconstitution of a functional human immune system in mice with severe combined immunodeficiency (SCID) [13]. Since then, the humanized mouse model has been improved to include various models with the immunodeficiency mouse strains. Many humanized mouse models have developed and widely been used for studies on cancer, infectious disease, transplantation or autoimmune disorders. Andrade *et al*. have successfully constructed reproducible systemic lupus erythematous (SLE)-humanized mouse model that produce human total-IgG and anti-double-stranded deoxyribonucleic acid Ab with PBMCs in SLE patients [14].

There are some standard approaches to develop human immune system into immunodeficient mice using PBMCs or hematopoietic stem cells (HSCs) in combination with implantation of autologous fragments of fetal thymus and liver [8,10,11]. The injection of human CD34^+^ HSC from bone marrow or cord blood into young mice is known as the Hu-RC-SCID model, that allows for the differentiation and development of immune system. Although the development of human T and B cells can be confirmed, insufficient maturation of these cells was observed [15,16]. The reason for insufficient maturation is that the human cell differentiation promoting factor is not found in the mouse cells, and human cells are refractory to mouse cell differentiation promoting factor [16]. Although humanized mouse models using HSC have been widely used, a humanized mouse model having sufficient anti-HLA Ab-producing ability corresponding to allo-antigen from PBMCs has not still been established. One of the reasons is that sufficient Ab production in the mouse body may not have been achieved even if the HSC engraftment was excellent, perhaps because human T and B cell interaction in the mouse spleen germinal center was insufficient. Another reason is that class switching from IgM to IgG is less likely to occur, the antigen-specific Ab producing ability is extremely low, and the Ab production promoting factor in the mouse body is refractory to human B cells, as in the previously reported humanized mice. Also, it is considered that the differentiation potential of B cell is low [16]. Humanized mouse models using mature PBMC but not HSC are still frequently used in various studies. As shown in previous reports [15], mere administration of human PBMCs into immunodeficient mice did not result in anti-HLA Abs production. Thus, we hypothesize that human immune cells may not be in sufficient contact with each other in the mouse, and that allo-antigen stimulation may not be transmitted to the various responder immune cells including human T, B, and antigen presenting cells in the mouse spleen. Given that *in vitro* mixed lymphocyte culture sufficiently transmits allo-antigen stimulation to the antigen presenting cells of responders, we speculated that transferring human responder matured and activated immunocompetent cells into the humanized mouse body after stimulating allo-antigen *in vitro* culture could produce allo-specific anti-HLA Abs.

To this end, two types of human PBMCs were co-cultured *in vitro* with human B cell differentiation promoting factor before injecting into naïve NSG mice. Therefore, we considered that CD40-CD40L signaling is required as an Ab-producing B cell activating factor. Ligation of CD40 on B cells by the CD40 ligand (CD154) on T cells promotes B cell proliferation, immunoglobulin production, isotype switching, and memory B cell generation [17]. The antigen-presenting cells (macrophage or dendritic cells), which recognize an antigen, present the antigen to the T cells, and Ab production occurs by the interaction between the T cells and B cells (CD40 and CD40L) [18-20]. We anticipated that antigen-specific Ab production might be enhanced by forcibly inducing the interaction via CD40 on the responder B cells by using feeder cells expressing h-CD40L. However, allo-specific anti-HLA Abs were not sufficiently detected by Ab production stimulating of CD40L alone, as shown in Figure 3. There are reports that BAFF-BAFF receptor signaling system, in addition to CD40-CD40L signaling, is essential for mouse B cell differentiation and proliferation [21,22]. As showing in the Figure 5 and 6, the use of feeder cells expressing both h-CD40L and h-BAFF did not increase human total-IgG and whole anti-HLA Abs. Even the induction of two typical signals that activate Ab-producing B cells resulted in sufficient anti-HLA Abs but suppressed production of allo-specific anti-HLA Abs. As an opposing idea, we attempted removing regulatory T cells, which negatively control a function of Ab-secreting B cells. We suspected that the responder T-regs might only suppress the activation of antigen-specific B cells. Although an alternate B cell activation signal was added or the T-regs that negatively control Ab-producing cells removed, non-specific anti-HLA Ab production was sufficient, but antigen-specific anti-HLA Ab production remain suppressed.

These results suggest that these humanized mouse models have a mechanism of allogeneic antigen-specific Abs production tolerance that is independent of the effects of h-BAFF-BAFF receptor signaling or T-reg regulation. This humanized mouse model is characterized by using human matured PBMCs from healthy volunteers, but not immature HSCs. Unlike the humanized mouse model administered with HSCs, that with PBMCs has the disadvantage of a shorter engraftment period that cannot be observed for a long time. However, since human PBMCs with a potential of producing some Abs are administered to mice under condition of Ab-producing B cell activation in vitro, the desired Ab production can be sufficiently identified within a relatively short period of 14-28 days. In future analyses of this humanized mouse model, PBMCs from clinical specimens will used, it might be considered as a simple and clinically applicable humanized model.

Interestingly, we have produced sufficient amounts of non-specific HLA Abs in the humanized mice, but not allo-specific anti-HLA Abs. We supposed that this may be due to induction of allo-antigen-specific tolerance during the *in vitro* culture with h-CD40L expressing mouse feeder cells, or B and T cells that respond to allo-antigen were exhausted and did not reach allo-specific anti-HLA Ab production enough. Also, we guessed that B and T cell clones that respond to allo-antigen in the *in vitro* mixed lymphocyte reaction may have been removed or induce anergy during the culture, developing a humanized mouse model in which allo-specific anti-HLA Abs are not produced.

We had initially tried to establish a humanized mouse model that produces allo-specific anti-HLA Abs. Conversely, a tolerant humanized model that produce non-specific anti-HLA Abs but not allo-specific was established. From these results, this anti-HLA Ab-producing humanized mouse model might be useful as a novel and profound model that can induce only antigen-specific B cells to tolerance.

## Conclusions

We attempted to develop allo-specific anti-HLA Abs-producing mice using humanized mice conditioned with human PBMCs. As a result of culturing feeder cells expressing both h-CD40L and h-BAFF for the purpose of activating Ab-producing B cells, sufficient non-specific anti-HLA Abs production was obtained, whereas allo-specific anti-HLA Abs production was unexpectedly suppressed. The availability and presence of a mechanism to negatively control the production of antigen-specific Abs in the *in vitro* culture system might make this anti-HLA Ab-producing humanized mouse a useful and profound model for the induction of tolerance of antigen-specific B cells. However, further investigation to elucidate the allogenic-specific tolerance mechanism in the modified humanized mouse model are needed.

## Acknowledgements

This work was performed in part at the Research Facilities for Laboratory Animal Science and the Analysis Center of Life Science, Natural Science Center for Basic Research and Development (N-BARD), Hiroshima University, Hiroshima, Japan; we thank the staff for their assistance.

We would like to thank Editage (www.editage.jp) for English language editing.

## Supporting information

**S1 Fig. Two types of PBMCs with different HLA types (combination 1) were administered to NSG mouse**.

In each mouse (# S-1 and S-2), only those on the responder side was detected in the peripheral blood of humanized mouse, and most h-CD45^+^ cells were h-CD3^+^ in FCM. The responder side is surrounded by a square.

**S2 Fig. Human total-IgG in mouse sera**

In each mouse, human total-IgG production was negligible.

**S3 Fig. Anti-HLA Ab titer in mouse sera**

Both of mice (#S-1, #S-2) could not produce anti-HLA Ab at all.

**S4 Fig. Expression of h-BAFF in FCM**

After transfection into the h-CD40L expressing NIH 3T3 fibroblasts, the rate of h-BAFF expression was over 90 %.

**S5 Fig. The ratio before and after T-regs removal in FCM**

The proportion of T-regs in PBMC was 16.2% (A) and decreased to 1.2 % after removal of T-regs (B) in FCM.

